# Distinct cortical spatial representations learned along disparate visual pathways

**DOI:** 10.1101/2024.10.10.617687

**Authors:** Yanbo Lian, Patrick A. LaChance, Samantha Malmberg, Michael E. Hasselmo, Anthony N. Burkitt

**Affiliations:** Department of Biomedical Engineering, The University of Melbourne, Melbourne, VIC 3010, Australia; Center for Systems Neuroscience, Department of Psychological and Brain Sciences, Boston University, Boston, MA 02215, USA; Graeme Clark Institute for Biomedical Engineering, University of Melbourne, VIC 3010, Australia

## Abstract

Recent experimental studies have discovered diverse spatial properties, such as head direction tuning and egocentric tuning, of neurons in the postrhinal cortex (POR) and revealed how the POR spatial representation is distinct from the retrosplenial cortex (RSC). However, how these spatial properties of POR neurons emerge is unknown, and the cause of distinct cortical spatial representations is also unclear. Here, we build a learning model of POR based on the pathway from the superior colliculus (SC) that has been shown to have motion processing within the visual input. Our designed SC-POR model demonstrates that diverse spatial properties of POR neurons can emerge from a learning process based on visual input that incorporates motion processing. Moreover, combining SC-POR model with our previously proposed V1-RSC model, we show that distinct cortical spatial representations in POR and RSC can be learnt along disparate visual pathways (originating in SC and V1), suggesting that the varying features encoded in different visual pathways contribute to the distinct spatial properties in downstream cortical areas.

**Conflict of interest statement:** The authors declare no competing financial interests.

## 1 Introduction

Animals perform very complex spatial navigation tasks, but how the brain’s navigational system processes spatial stimuli to guide behavior is still unclear. In recent decades, experimental studies of brain navigation have identified many different types of spatial cells, including place cells (O’Keefe and Dostrovsky, 1971; O’Keefe, 1976), head direction cells (Taube et al., 1990a,b), grid cells (Hafting et al., 2005; Stensola et al., 2012), boundary cells (Solstad et al., 2008; Lever et al., 2009) and speed cells (Kropff et al., 2015; Hinman et al., 2016). Many of these cells code for an allocentric spatial map, which is defined with respect to the external environment. Recently, increasingly more experimental studies in the rodent brain have uncovered spatial cells that are egocentric, or defined with respect to the animal itself, in different brain areas, including lateral entorhinal cortex (Wang et al., 2018), dorsal striatum (Hinman et al., 2019), postrhinal cortex (POR) (Gofman et al., 2019; LaChance et al., 2019; LaChance and Taube, 2023; LaChance and Hasselmo, 2024), and the retrosplenial cortex (RSC) (Alexander et al., 2020; LaChance and Hasselmo, 2024).

Alexander et al. (2020) identified egocentric spatial cells in RSC that are selective to the boundaries of the arena with a preferred self-centered orientation (e.g., left, right, front or back) at a preferred distance, with the population of RSC egocentric cells displaying a distribution of preferred orientations and distances. In an experimental study of neurons in the rat POR, LaChance et al. (2019) discovered egocentric spatial cells that encode the egocentric bearing and distance of the geometric center of a square arena. In a follow-up study investigating the rat POR in square and L-shape arenas, LaChance and Taube (2023) found that POR egocentric cells can encode both local and global aspects of environmental geometry. Moreover, recent work by LaChance and Hasselmo (2024) showed that RSC and POR have distinct codes for environment structure and symmetry by simultaneously recording cells from both brain areas.

Animals use their sensory system, that is egocentric in nature, to explore their spatial environment. Consequently, understanding how an egocentric representation of space arises from sensory input during learning is vital to understanding the brain’s navigational system. Using a neural network model with synaptic plasticity, our previous work showed that egocentric cells in RSC can be learnt from the visual input of the primary visual cortex (V1), which can also account for the diversity of RSC cell properties (Lian et al., 2023).

Nevertheless, how POR egocentric spatial cells develop via a learning process is still unknown, and the underlying mechanisms that lead to distinct egocentric spatial codes in RSC and POR remain to be understood. Solving these important problems will help us better understand how the brain’s navigational system develops, and it will assist in identifying the underlying principles of neural mechanisms subserving navigation.

Although both RSC and POR receive visual sensory input, the visual information they receive comes from distinct pathways: RSC primarily receives visual input from V1 (van Groen and Wyss, 1992) while POR primarily receives visual information via the superior colliculus (SC) (Zhou et al., 2018; Bennett et al., 2019; Beltramo and Scanziani, 2019; Brenner et al., 2023). Moreover, cells in V1 and SC have different functional properties. Cells in V1, such as simple and complex cells, are selective to bar-like features (Carandini, 2006), while SC cells are selective to motion contained in the visual input that may reflect processing of optic flow information (Ahmadlou and Heimel, 2015; Li et al., 2020; Ge et al., 2021; Teh et al., 2023).

In this study, we build and investigate a neural network learning model of POR cells based on the SC to POR pathway. As a virtual rat runs freely in a simulated environment, the visual input of the virtual rat is captured and then used as the input to train the neural model in which the neural connections between SC and POR are updated according to the activity-driven synaptic plasticity of the model. Our results demonstrate that this model learns various types of POR egocentric spatial responses that have been observed in experiments (LaChance et al., 2019; LaChance and Taube, 2023). Additionally, combining our previous model of RSC cells based on the V1 to RSC pathway, these two models can account for the distinct egocentric spatial codes found in a recent experimental study (LaChance and Hasselmo, 2024). Our study illustrates how POR egocentric spatial cell properties can be learnt from visual input via the SC pathway, and it indicates how different visual processing mechanisms in V1 and SC could be the origin of distinct egocentric spatial codes in RSC and POR, respectively. Our models are based on the principle of sparse coding, indicating that sparse coding may be one of the fundamental principles of the brain’s navigational system.

## 2 Methods

### 2.1 The simulated environments, trajectory and visual input

#### 2.1.1 Environments

The simulated environments were created to mimic the environments in the recent experimental study of POR and RSC (LaChance and Hasselmo, 2024), including a square arena with one or two white cue cards on walls and a L-shape arena with one white cue card on one wall.

#### 2.1.2 Trajectory

Similar to the study by D’Albis and Kempter (2017), the running trajectory ***r***_*t*_ is generated from the stochastic process described by the equation:

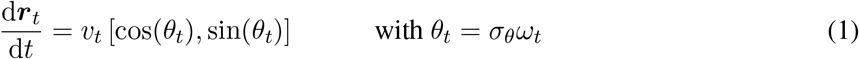

where *v*_*t*_ is the speed sampled from an Ornstein-Uhlenbeck process with long-term mean 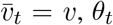 is the direction of movement, *ω*_*t*_ is a standard Wiener process, and *σ*_*θ*_ is the parameter that controls the tortuosity of the running trajectory. When the virtual rat is running toward the wall and very close to the wall (within cm), the running direction of the rat (*θ*_*t*_) is set to the direction parallel to the wall. If the rat location generated by Eq. 1 falls outside of the environment, the stochastic process generates alternative iterations until a valid location is generated. The running trajectory of the virtual rat is generated at 20 Hz; i.e., the position is updated every 50 ms according to Eq. 1. The long-term mean speed, *v*, is set to 30 cm/s. For each session, the virtual rat runs for 1800 s.

#### 2.1.3 Visual input

We use the Panda3D game engine (panda3d.org), an open-source framework for creating virtual visual environments, to create the environments and generate the corresponding visual input of the virtual rat along the trajectories generated above. The visual input of the simulated animal is modelled using a camera with a 150° field of horizontal view to mimic the wide visual field of rat and a 90° field of vertical view. The visual input at each time stamp is a 150*×*90 pixel image where each pixel represents one degree of the visual field. The camera is always facing the front, meaning that the head direction is aligned with the movement direction for the simulated animal.

### 2.2 Learning egocentric cells in POR

In this study, our model of learning of the response properties of POR egocentric spatial cells is based on the experimental evidence that POR receives visual information primarily via the superior colliculus (Beltramo and Scanziani, 2019; Brenner et al., 2023).

#### 2.2.1 Vision processing in superior colliculus

Cells in the superior colliculus (SC) respond to motion contained in the visual input and the preferred motion direction of each cell depends on its position in the visual field (Li et al., 2020). Specifically, the global map of visual motion selectivity of SC neurons is bilaterally symmetric and is biased towards upward motion, as seen in (Li et al., 2020, Fig. 7). In this study, we create a global map of SC neuron visual motion selectivity and build a mathematical model for SC neurons whose responses depend on motion speed and direction. The detailed process is as follows: First, the 150*×*90 visual input is down-sampled by a factor of 5, which reduces to 30*×*18. Second, the 30*×*18 visual input is used to compute the optic flow; at each point of the 30*×*18 grid, the corresponding optic flow has a motion direction and motion speed. Third, a global map of preferred motion direction is created for model SC neurons, as shown in Fig. 1. Fourth, for each SC neuron, the response is computed based on its position in the visual field, preferred motion direction, and preferred motion speed. At each position of the visual field, there are five different speed preferences (2, 4, 8, 16, or 32 degrees per second) and four different direction preferences uniformly sampled between *θ*_map_ − 30° and *θ*_map_ + 30° where *θ*_map_ is the direction determined by the global motion direction map, illustrated in Fig. 1. Therefore, there are altogether 10,800 (=30*×*18*×*5*×*4) SC neurons in the model.

**Figure 1.**
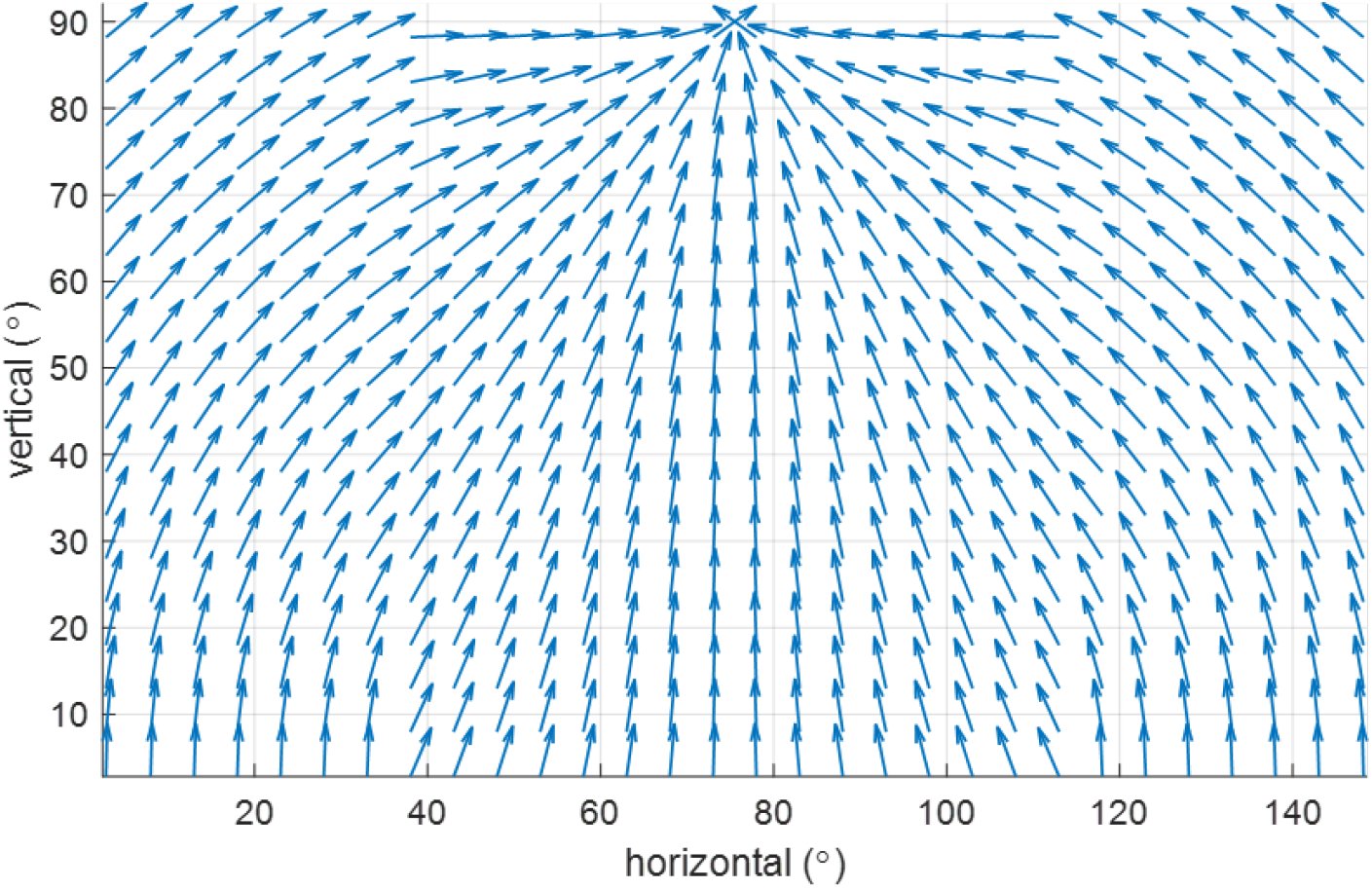
The global map of preferred motion direction of vision processing in superior colliculus (SC). The preferred direction of one SC visual cell depend on its position in the visual field. In general, SC visual cells prefer upward motion relative to the central vertical axis of the visual field. This figure is adapted from (Li et al., 2020, Fig. 7).

The response of each SC neuron at any position is given by

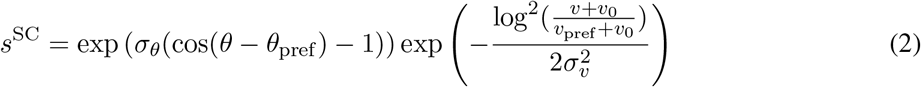

where *θ* is the direction of the optic flow, *θ*_pref_ is the preferred motion direction, *σ*_*θ*_ = 1.5 is the bandwidth of direction selectivity, *v* is the speed of the optic flow, *v*_pref_ is the preferred speed, *v*_0_ = 0.33 is the speed offset, and *σ*_*p*_ = 1.16 is the bandwidth of speed selectivity (Beyeler et al., 2016).

#### 2.2.2 SC-POR model: modelling POR cells using visual input from SC

Our previous learning model of RSC egocentric boundary cells is built on the anatomical connection from V1 to RSC, as illustrated in Fig. 2B, and the proposed V1-RSC model applies the principle of sparse coding to the input from V1 cells that process the visual input according to their preferred spatial features (Lian et al., 2023). Similar to the structure of V1-RSC model (Lian et al., 2023), we build a SC-POR model, illustrated in Fig. 2A, based on the visual pathway from SC to POR (Brenner et al., 2023) using the principle of sparse coding that takes the SC responses as the input.

**Figure 2.**
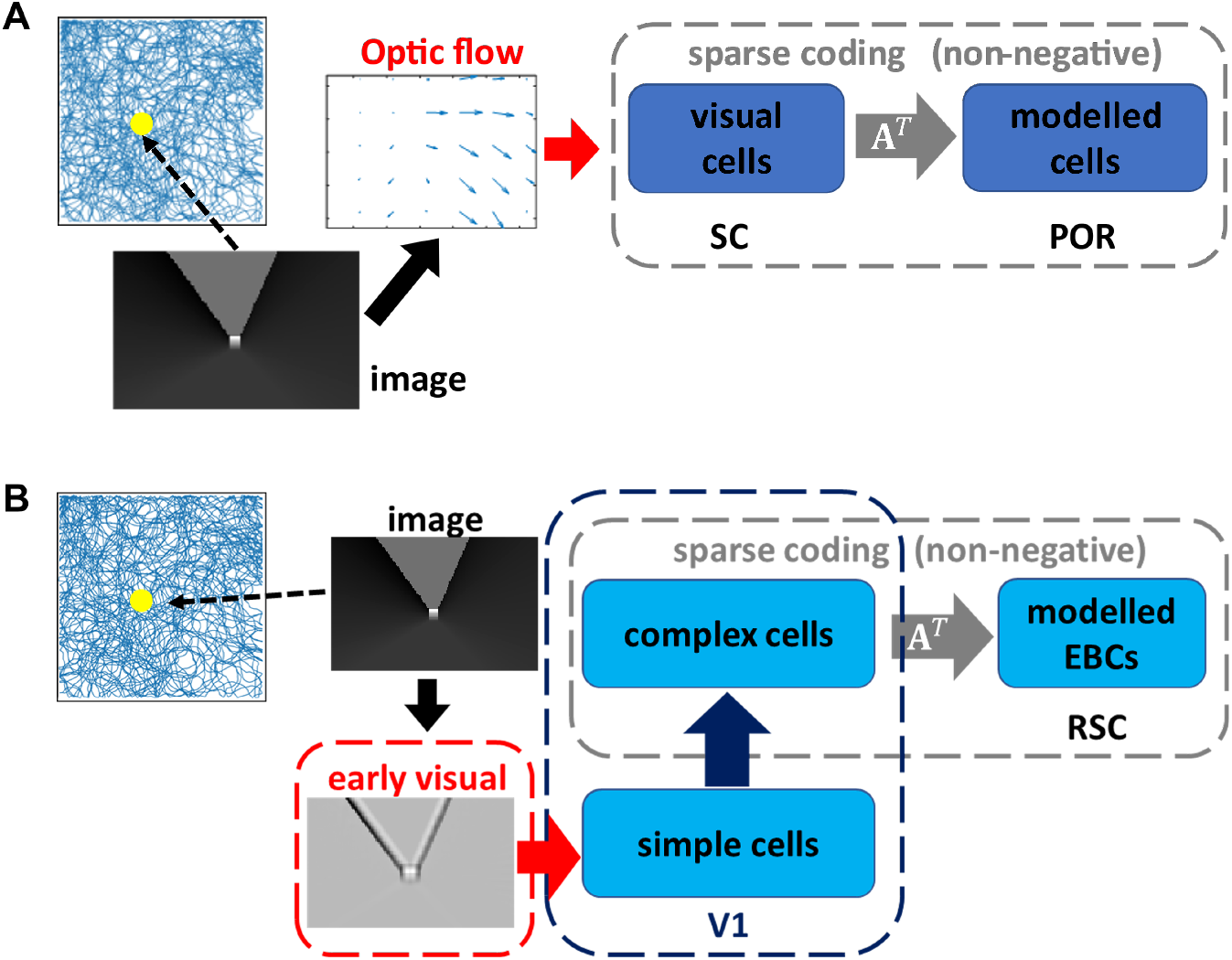
Structures of SC-POR model and the previously developed V1-RSC model. The simulated animal runs in the trajectory (see Section 2.1.2) in the simulated environment. The simulated visual scene the animal sees at different locations is the visual stimulus of the simulated animal. A) SC-POR model: the optic flow is computed from the raw visual input and then used to generate responses of SC visual cells that are selective to different motion speeds and directions; SC cells then project to modelled POR cells and a SC-POR network is implemented based on non-negative sparse coding. B) V1-RSC model: the raw visual input is pre-processed by the early visual system and then projected to V1 that involves simple cell and then complex cell processing; complex cells in V1 then project to modelled RSC cells and a V1-RSC network is implemented based on non-negative sparse coding (Lian et al., 2023).

In this study we implemented both the V1-RSC and SC-POR models, and the V1-RSC model is exactly the same as in our previous study (Lian et al., 2023). The major difference between the previously proposed V1-RSC model and the new SC-POR model presented here is the response of model visual cells. For the V1-RSC model, the images captured by the camera of the virtual rat undergo feature selection processing in V1 and the response of model V1 cells has no movement dependence. However, for the SC-POR model, the response of model SC cells has movement dependence because of the properties described in Section 2.2.1.

#### 2.2.3 Implementing SC-POR model and V1-RSC model

The model dynamics and learning rule of implementing SC-POR model is similar to our previous study of the V1-RSC model (Lian et al., 2023), as described by

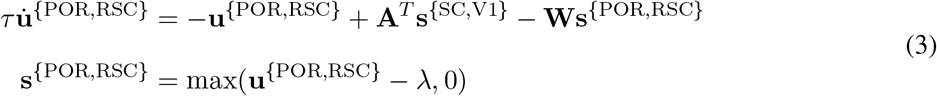

And

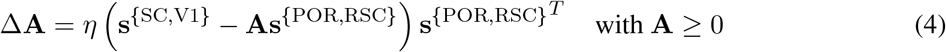

where **s**^{SC,V1}^ is the visual input of cells in SC (Eq. 2 and Fig. 2A) or V1 complex cells (Fig. 2B), **s**^{POR,RSC}^ represent the response (firing rate) of the model neurons in the POR or RSC, **u**^{POR,RSC}^ can be interpreted as the corresponding membrane potential, **A** is the matrix that represents the connection weights between SC visual cells and model neurons in the POR (Fig. 2A) or between V1 cells and model neurons in the RSC (Fig. 2B), **W** = **A**^*T*^ **A** − 1 and can be interpreted as the recurrent connection between model neurons in the POR or RSC, 1 is the identity matrix, *τ* is the time constant of the model neurons in the RSC, *λ* is the positive sparsity constant that controls the threshold of firing, and *η* is the learning rate. Each column of **A** is normalised to have length 1. Non-negativity of both **s**^{POR,RSC}^ and **A** in Eqs. 3 & 4 is incorporated to implement non-negative sparse coding.

##### Training

The training of the models is as follows: For the implementation of SC-POR and V1-RSC models, there are 100 model neurons in POR or RSC and the parameters are given below. For the model dynamics and learning rule described in Equations 3 & 4, *τ* is 10 ms, the time step of implementing the model dynamics is 0.5 ms. The simulated visual input generated at different positions along the simulated trajectory is used to train the model. Since the simulated trajectory is generated at 20 Hz, at each position of the trajectory, there are 100 iterations of computing the model response using Equation 3. After these 100 iterations, the learning rule in Equation 4 is applied such that connection **A** is updated. The animal then moves to the next position of the simulated trajectory. As the inputs to the model, **s**^SC^ and **s**^V1^, have different statistics, some hyperparameters are slightly different when training the model. For SC-POR model, *λ* is set to 1 and the learning rate *η* is set to 0.1, while *λ* is set to 0.2 and the learning rate *η* is set to 0.3 for V1-RSC model. For both models, the learning rate *η* is set to be 10% of the original value for the final 15% of the simulated trajectory.

### 2.3 Data Collection

#### 2.3.1 Recording environment and manipulations

The experimental environment consisted of a 1.2 × 1.2 m square box with 60 cm high walls. The walls and floor were painted black, and a large white cue card (cue A) with a width of 72 cm was placed along the south wall and covered the full vertical extent of the wall. A black floor-to-ceiling curtain surrounded the environment to block visual perception by the animals of global room cues. Baseline recording sessions involved animals foraging for randomly scattered sugar pellets in this environment for 20 min.

To examine the tuning of egocentric bearing (EB) cells to local or global aspects of environmental geometry, local and global geometric cues were placed into conflict by adding additional walls into the environment to block access to the northeast quadrant, transforming the environment into an L-shape. Recording sessions in the L-shape lasted 15-20 min.

To examine the responses of head direction (HD) cells to imposed symmetry of visual landmarks, we sometimes followed the baseline session with a second 20 min session where an identical cue card was placed along the north wall (cue B), making the environment visually symmetrical. Bidirectional responses of HD cells were then assessed by comparing cell responses in the initial session with cue A only (A1 session) to the session with both cues (AB session).

#### 2.3.2 Experimental data collection

To assess the similarity of modeled cells to experimentally recorded cells, we used a previously published dataset of neurons recorded from the POR and RSC of female rats during random foraging in an open field environment that was manipulated to test specific properties of EB and HD cells (LaChance and Hasselmo, 2024). These same experiments and relevant analyses were simulated in the current study. Methods concerning electrophysiological data acquisition can be found in (LaChance and Hasselmo, 2024), while methods concerning behavior and data analysis used for both experimental and model cells are included here.

#### 2.3.3 Model data collection

For each of the SC-POR and V1-RSC models, we train the model in the 1.2 × 1.2 m square arena with one white cue card (A1 session) to mimic the baseline session in the experimental study (LaChance and Hasselmo, 2024). In this session, both models undergo a learning process according to the procedure described in previous sections. After the models finish learning, the models are tested (i.e., no learning with *η* = 0) in different environments (A1/baseline session, L-shape session, AB session) to collect model responses for further analysis. Both models are rate-based descriptions of the neural activity, so model responses are then transformed into spikes using a Poisson spike generator with a maximum firing rate 30 Hz for the whole modelled population.

### 2.4 Data analysis

#### 2.4.1 Cell classification with a generalized linear model

Both experimental and model cells were classified as encoding one or more behavioral variables using ten-fold cross-validation with a Poisson Generalized Linear Model (GLM), as used in previous studies (Hardcastle et al., 2017; LaChance et al., 2019; LaChance and Hasselmo, 2024). Experimental cells were classified as encoding one of four behavioral variables: egocentric bearing of the environment center (proxy for tuning to outer boundaries); egocentric distance of the environment center; allocentric head direction; and linear speed. Linear speed was omitted from the classification procedure for model cells as the simulated trajectory maintained a relatively constant speed.

Use of this classification scheme has been described previously (LaChance et al., 2019; LaChance and Hasselmo, 2024). Briefly, the spike train of a given cell was estimated as the firing rate vector *r* using the following equation:

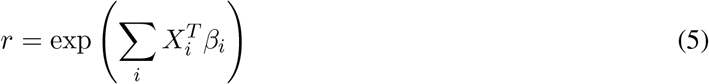

where *X* is a design matrix containing animal state vectors for a given behavioral variable across time points *T, β* is the estimated parameter vector for that variable, and *i* indexes across variables included in the model. The estimated parameter vectors were optimized by maximizing the log-likelihood *l* of the real spike train *n* across time points *t* using the following equation:

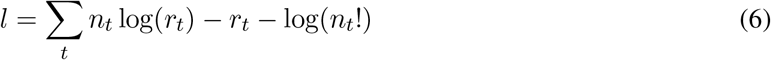

A smoothing penalty was also incorporated to avoid overfitting, which enforced minimal differences between adjacent bins of each parameter vector:

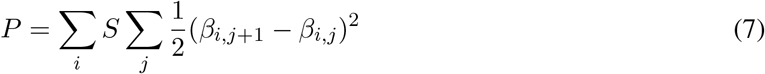

where *S* is a smoothing hyperparameter (20 for all variables), *i* indexes across variables, and *j* indexes across elements in the parameter vector for each variable. SciPy’s *optimize*.*minimize* function was used to estimate response parameters by minimizing (*P* − *l*). Thirty bins were used for egocentric bearing and allocentric head direction parameter vectors, and ten bins were used for egocentric distance and linear speed.

For each fold of the cross-validation procedure, the recording session was split into training (9/10) and testing (1/10) data. The full model containing all variables was first optimized on the training data, from which the parameter estimates were extracted and used to create all possible smaller models, which were evaluated on the testing data using log-likelihood increase relative to an intercept-only model. This procedure occurred 10 times, until all parts of the data had been used as testing data.

For model selection, a forward search procedure was used (Hardcastle et al., 2017). Briefly, the loglikelihood values from the best one-variable model were compared to the log-likelihood values from the best two-variable model that contained that variable using a one-sided Wilcoxon signed-rank test. If the two-variable model performed significantly better, it was compared to the best three-variable model that contained those two variables, and so on. If the more complex model did not perform significantly better, the simpler model was chosen. The final chosen model was then compared to an intercept-only model that only contained the cell’s mean firing rate, and if it performed significantly better than the intercept-only model, it was chosen as the cell’s classification. Otherwise, the cell was considered ‘unclassified’.

#### 2.4.2 Tuning curves and final cell classifications

For EB and HD measurements, tuning curves were created for each using 12° bins. For each cell, a tuning curve was constructed by dividing the number of spikes associated with each bin by the amount of time the animal spent occupying that bin. The Mean Vector Length (MVL) and mean angle of that tuning curve were used to assess the cell’s tuning strength and preferred direction, respectively. A cell was considered an EB or HD cell if it: i) passed the GLM classification procedure for EB or HD tuning (discussed above); ii) had an MVL that passed the 99^th^ percentile of a within-cell shuffle distribution (discussed below); iii) had a peak firing rate that exceeded 1 Hz. A hard MVL cutoff was also imposed (0.10 for EB cells and 0.15 for HD cells).

#### 2.4.3 Local vs. global GLM

To test if individual cells were more strongly tuned to local geometric features or the global structure of the environment in the square and L-shaped arenas, a Poisson GLM was used to compare between these two possibilities. The global version of the model was identical to the classification GLM (without the smoothing component), and included the following variables: egocentric bearing of the environment center, egocentric distance of the environment center, allocentric head direction, and linear speed (for experimental cells only). Tuning to the environment centroid is mathematically equivalent to tuning to the full extended boundary of the environment (i.e., global geometry tuning).

In contrast, the local version of the model replaced the egocentric bearing and distance of the centroid with the egocentric bearing and distance of the nearest two walls. To accomplish this, for each time point in the recording session we calculated the egocentric bearing and distance of the closest point along each of the nearest two walls. For each wall, two animal state vectors were created, 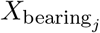 and 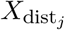, where *j* indicates measurements made relative to the *j*-th closest wall. We then solved for the optimal parameter vectors *β*_bearing_ and *β*_dist_ (along with HD and speed parameters) by optimizing the GLM as described above, this time modeling the cell’s firing rate as:

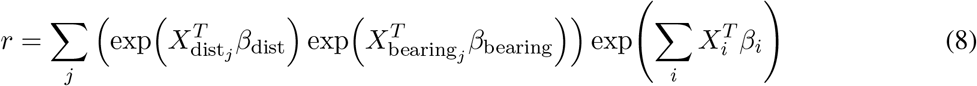

such that the cell’s response to the bearing of each wall is scaled by the distance of each wall and then summed before being multiplied by the responses to other behavioral variables (HD and linear speed; indexed by *i*). Because we were interested in the explanatory power of each model, we omitted the smoothing component and trained and tested the models on the full recording session. We then computed a Globality Index (GI) that compared the log-likelihood fits of the local and global models relative to a uniform model that only included the cell’s mean firing rate:

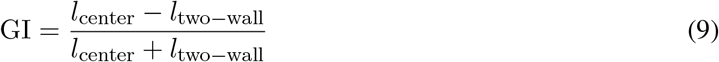

Values of GI could potentially range from -1 (strictly local) to +1 (strictly global), although due to collinearity of center and wall measurements, they are generally closer to 0.

#### 2.4.4 Four-fold symmetry analyses

To assess four-fold symmetry of EB cell firing in the square environment, we assessed symmetry based on three different assumptions for cells with four-fold responses:

1. HD tuning curves. Cells should preferentially respond to four distinct allocentric HDs, such that their HD tuning curves possess four distinct peaks spaced 90° apart. To assess this property, we created an autocorrelation function for each cell’s HD tuning curve by correlating the original curve with a shifted version at all possible directional offsets (i.e., across all 12° bins of the tuning curve). This autocorrelation function was used to compute a symmetry score (described below).
2. HD x location correlation structure. Cells should have four distinct firing fields that are each associated with a different HD. To assess this property, we created separate firing rate maps for time periods when the animal was facing separate HDs. Rate maps were created for HDs from 0° to 360° in 3° increments, with each rate map consisting only of times points that the animal faced that particular HD (*±*30°). We then computed correlations between all possible pairs of rate maps in order to produce a correlation matrix. Cells with four discrete firing fields associated with four discrete HDs should show four discrete ‘blocks’ of high correlation value along the main diagonal of the matrix, each with a width of approximately 90°. An autocorrelation function was computed for the central 90° of this matrix by shifting it along its main diagonal. This autocorrelation function was used to compute symmetry scores (described below).
3. Radial symmetry of firing fields. The cell’s HD-associated firing fields should be systematically placed radially at 90° offsets relative to the center of the environment. To assess this property, we used a GLM to model each EB cell’s spike train using 1-dimensional (1D) distance and rotation functions (in addition to allocentric HD). The distance function was projected across the environment to create a pseudo-2D rate map, and could be rotated about the environment center according to the animal’s HD. The degree of rotation associated with each HD was determined by the rotation function, which should have a ‘stepwise’ appearance for cells with four-fold symmetry (with steps spaced 90° apart), as the cells ‘snap’ to a new firing field every 90° of rotation. The GLM was optimized in the same way as the classification and local vs. global GLMs, but included an additional penalty imposed on the mean vector length of the rotation function to ensure sampling of the full range of possible rotations. Thirty bins were used for each variable. Following optimization, the rotation function was detrended by subtracting a linear range of angles from 0° to 360°, after which an autocorrelation was computed from the detrended function and symmetry scores were computed (described below).

#### 2.4.5 Computation of symmetry scores

To transform the 1D autocorrelations into four-fold symmetry scores, we took the lowest correlation value at 90°, 180°, or 270° and subtracted the highest correlation value at 45°, 135°, 225°, or 315°. This method is similar to the technique used to detect hexagonal symmetry of grid cell firing (Hafting et al., 2005).

#### 2.4.6 Allocentric location firing rate maps

To assess firing rate distributions over allocentric space, we divided the animal’s 2D location throughout the recording session into 4 cm × 4 cm bins. For each cell, the number of spikes associated with each bin was divided by the amount of time the animal spent occupying that bin. The resulting firing rate map was smoothed with a Gaussian filter.

#### 2.4.7 Place-by-HD vector plots

To visualize a cell’s HD preferences across allocentric space, we partitioned the environment into 8 × 8 spatial bins and created HD tuning curves based on the time points the animal spent occupying each bin. 30° bins were used, as the occupancy time for each bin tended to be small. If the occupancy for a bin was under 200 ms, the bin was expanded in steps of 1 cm until the 200 ms threshold was met, similar to ‘adaptive binning’ in previous studies (Skaggs et al., 1996; Wang et al., 2018). The MVL and preferred direction were computed for each bin and plotted as the length, 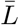, and direction, *θ*_pref_, of the resulting vector, respectively. Bins with peak firing rates smaller than 1 Hz were omitted.

#### 2.4.8 Assessment of HD cell bidirectionality

To determine if HD cells became bidirectionally tuned in the cue duplication experiment (i.e., fired in two opposite directions), we computed a Bidirectionality Index (LaChance et al., 2022; LaChance and Hasselmo, 2024). Two tuning curves were created for each HD cell: one based on the animal’s actual HD; and one where the animal’s HD had been doubled. Angle doubling can be used to transform a symmetrical bimodal distribution into a unidirectional one. The bidirectionality index (BI) was then calculated from the resulting MVLs, MVL_normal_ and MVL_doubled_, as follows:

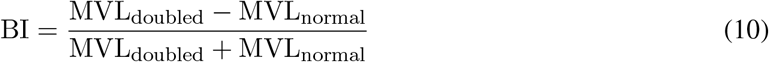

#### 2.4.9 Cue modulation measures

To assess the extent to which HD cell responses could attributed to each cue in the AB session, we fitted a bidirectional von Mises distribution (two peaks or troughs separated by 180°) to each cell’s HD tuning curve in the AB session (LaChance et al., 2022; LaChance and Hasselmo, 2024). As only POR and SC-POR cells tended to show trough-locked tuning (i.e., cells that are inhibited when the animal faces a certain direction), trough fits were only used for POR and SC-POR cells with maximal firing rates oriented away from the cue card, and RSC and V1-RSC cells were modeled using peak fits. Modulation by cue A was assessed by finding the von Mises peak or trough closest to the cell’s A1 peak or trough, then computing the firing rate difference between that peak or trough and the minimum or maximum of the fit curve, respectively. This firing rate difference was then transformed into a modulation index (MI) by dividing it by the maximum firing rate of the fit curve (fr = firing rate):

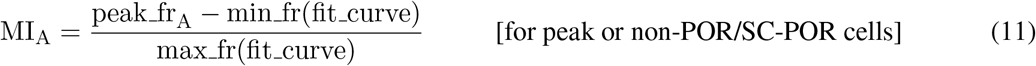

OR

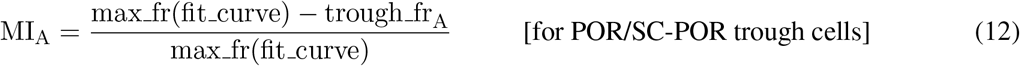

where A indicates the portion of the tuning curve associated with cue A. The MI for cue B was calculated by performing the same computation on the peak or trough 180° opposite.

#### 2.4.10 Shuffling procedure

Each cell’s spike train was randomly shifted by at least 30 s, with time points that extended beyond the end of the session wrapped to the beginning, in order to offset the spike data from the tracking data. Relevant tuning scores were computed based on the shifted spike train. This procedure occurred 400 times for each cell, and a 99^th^ percentile within-cell cutoff was used to determine significance of tuning for each cell.

#### 2.4.11 Statistics

All statistical tests were nonparametric and two-sided, except for GLM classification comparisons which were one-sided Hardcastle et al., 2017; LaChance et al., 2019) and used an level of 0.05. Paired comparisons were made using Wilcoxon signed rank tests, while unpaired comparisons used a rank-sum test.

## 3 Results

### 3.1 Firing properties of experimental and model egocentric bearing cells

Both SC-POR and V1-RSC models produced simulated neurons with robust spatial tuning. To classify the model cells, we used both tuning curve analyses and cross-validation with a generalized linear model (GLM) to confirm tuning to one or more of the following three spatial variables: 1) egocentric bearing (EB) of the environment boundaries/center; 2) egocentric distance (ED) of the environment boundaries/center; 3) allocentric head direction (HD; see Methods). Among the 100 SC-POR cells simulated in a 1.2 × 1.2 m square arena, 72% were classified as EB cells, 25% as ED cells, and 54% as HD cells. For the 100 V1-RSC cells, those numbers were 90%, 64%, and 62%, respectively, as shown in Fig. 3A. In both models, the cells often exhibited conjunctive tuning, such that many cells were tuned to more than one variable (66% of SC-POR cells and 82% of V1-RSC cells). While the overall percentage of cells that encoded at least one variable (SC-POR: 84%; V1-RSC: 93%) was higher than that observed in an experimental dataset (POR: 47%; RSC: 53%; LaChance and Hasselmo, 2024), the presence of EB, ED, and HD cells, including many conjunctive cells, matched well with the experimental data, as shown in Fig. 3A, C.

**Figure 3.**
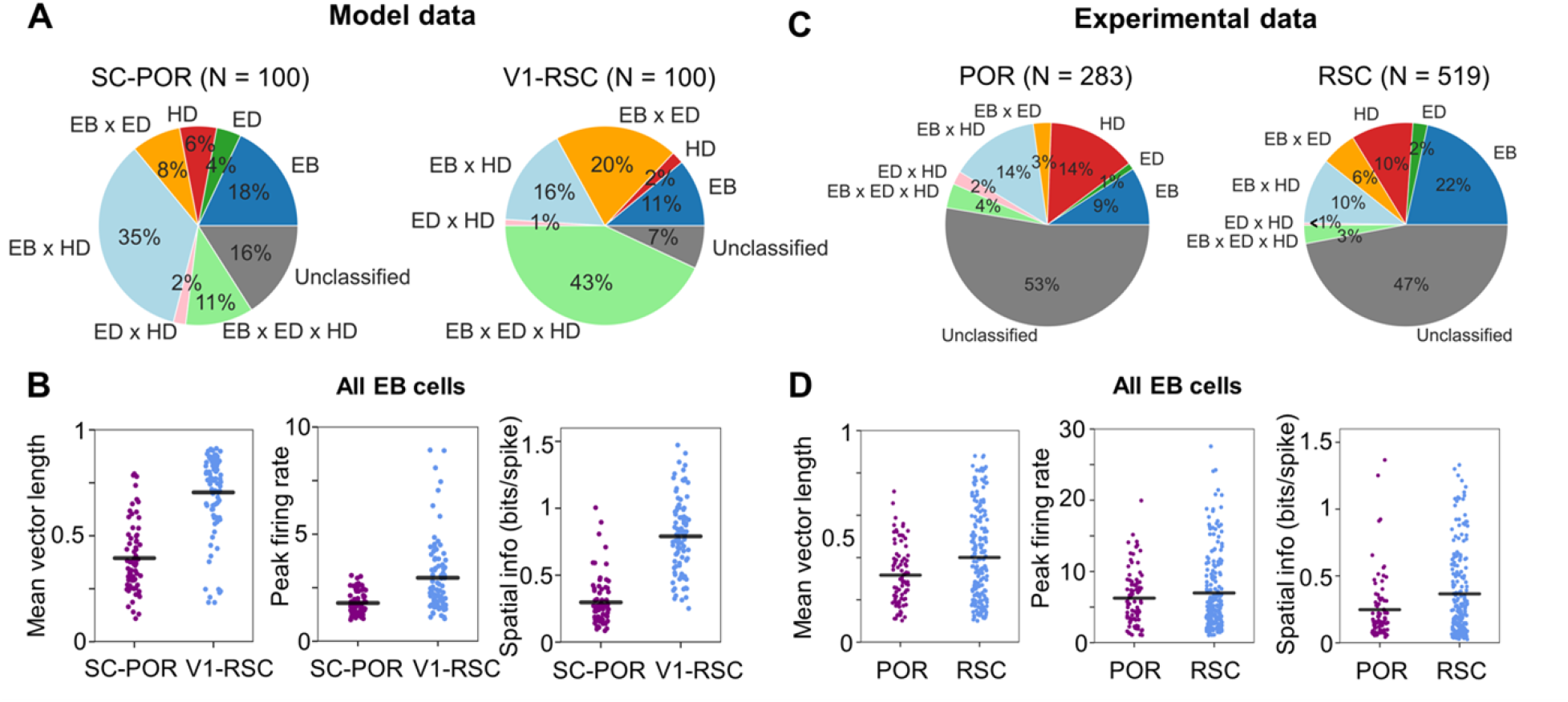
Classifications and egocentric bearing cell statistics. A) Percent of modeled cells classified as encoding one or more of three behavioral variables: egocentric bearing (EB), egocentric distance (ED), and allocentric head direction (HD). B) Population statistics of EB cells compared between SC-POR and V1-RSC models. From left to right: egocentric bearing mean vector length; peak firing rate; spatial information content. Note that V1-RSC cells tended to have higher values in all three domains. C-D) Same as (A-B) but for experimental neurons recorded from POR and RSC.

Focusing on EB cells, the baseline firing properties of the SC-POR and V1-RSC cells differed in a similar way to the experimental POR and RSC EB cells, as shown in Figs. 4-7, with V1-RSC cells generally exhibiting higher mean vector lengths (MVLs; Z = 7.96, P = 1.66e-15), peak firing rates (Z = 5.99, P = 2.10e-9), and spatial information content (Z = 9.89, P = 4.79e-23) than SC-POR cells, as shown in Fig. 3B. Overall, SC-POR cells tended to have broad tuning profiles with firing that covered a large portion of the environment, shown in Fig. 5, compared to V1-RSC cells that fired in a more restricted set of directions and locations, shown in Fig. 7. These trends were apparent in the experimental data (Figs. 4, 6), though measurement differences only reached significance for MVLs (Z = 2.55, P = 0.011) and spatial information content (Z = 2.16, P = 0.030) but not peak firing rates (Z = 0.11, P = 0.91, as shown in Fig. 3D.

**Figure 4.**
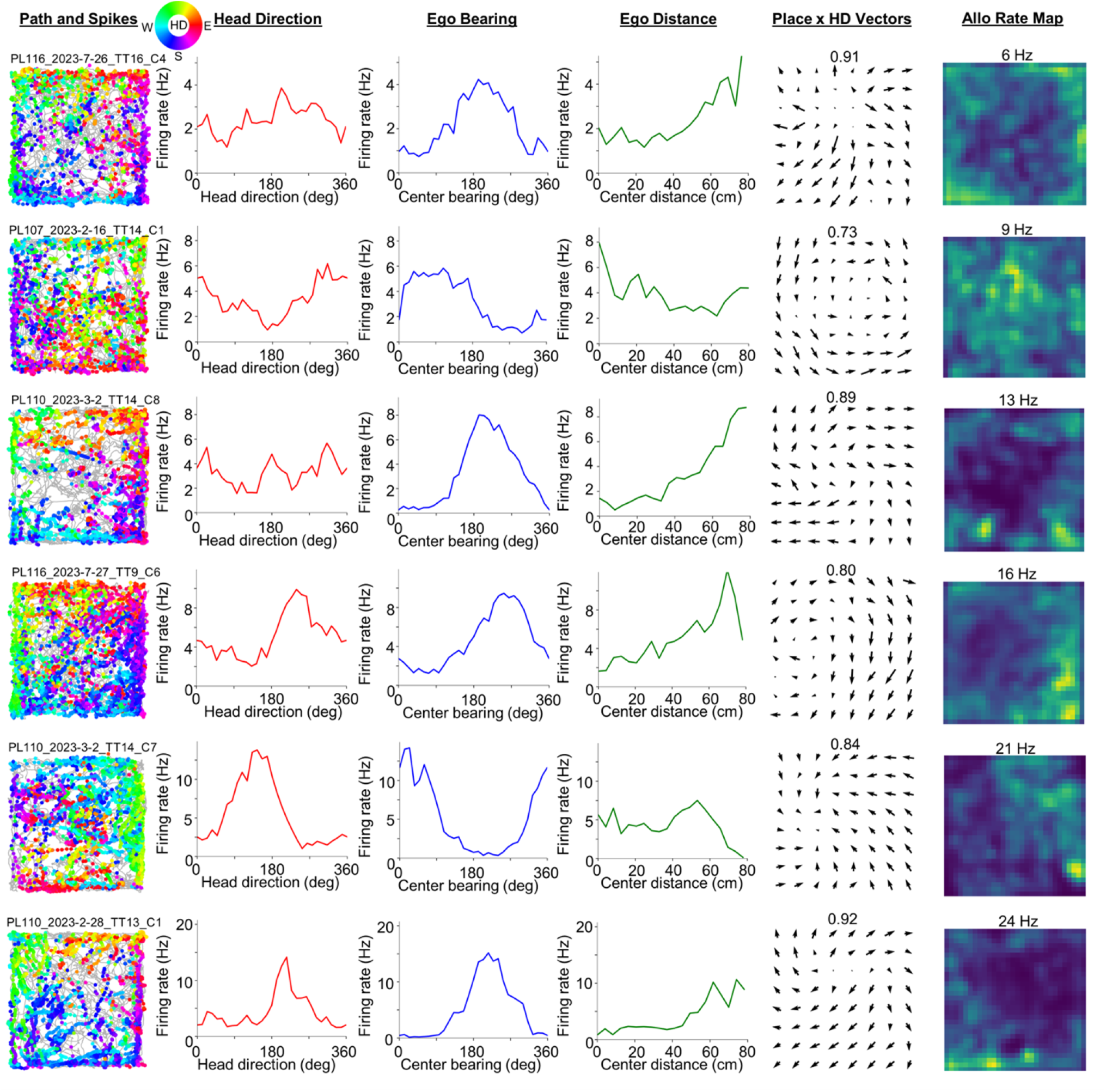
Experimental POR egocentric bearing cells. Directional spike plots, tuning curves, place-by-HD vector plots, and allocentric firing rate maps for six example experimental POR cells with significant egocentric bearing tuning. The number above the place-by-HD vector plot indicates the highest MVL, while the number above the allocentric rate map indicates the cell’s peak firing rate.

**Figure 5.**
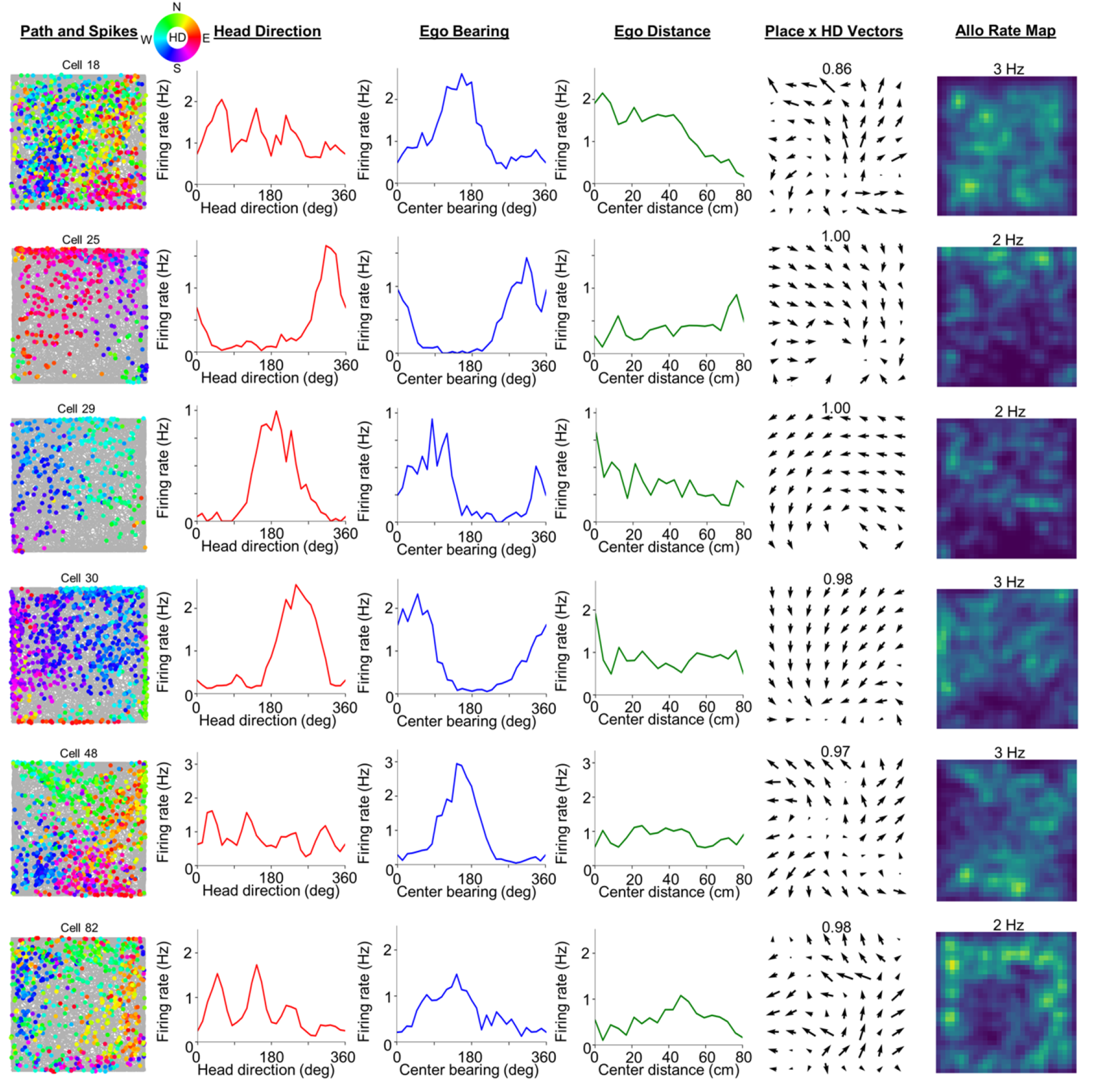
Modeled SC-POR egocentric bearing cells. Directional spike plots, tuning curves, place-by-HD vector plots, and allocentric firing rate maps for six example modeled SC-POR cells with significant egocentric bearing tuning. The number above the place-by-HD vector plot indicates the highest MVL, while the number above the allocentric rate map indicates the cell’s peak firing rate.

**Figure 6.**
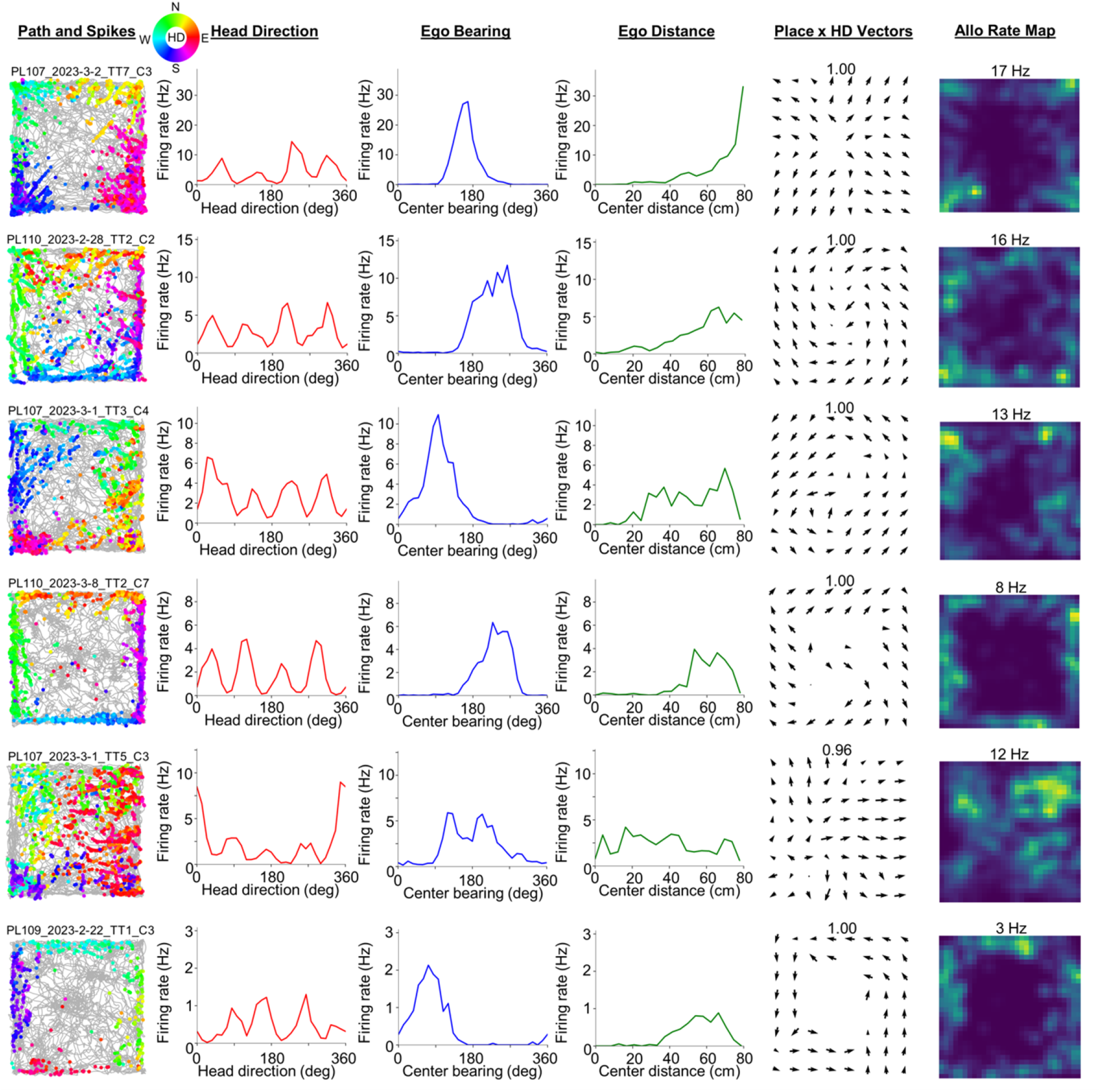
Experimental RSC egocentric bearing cells. Directional spike plots, tuning curves, place-by-HD vector plots, and allocentric firing rate maps for six example experimental RSC cells with significant egocentric bearing tuning. The number above the place-by-HD vector plot indicates the highest MVL, while the number above the allocentric rate map indicates the cell’s peak firing rate.

**Figure 7.**
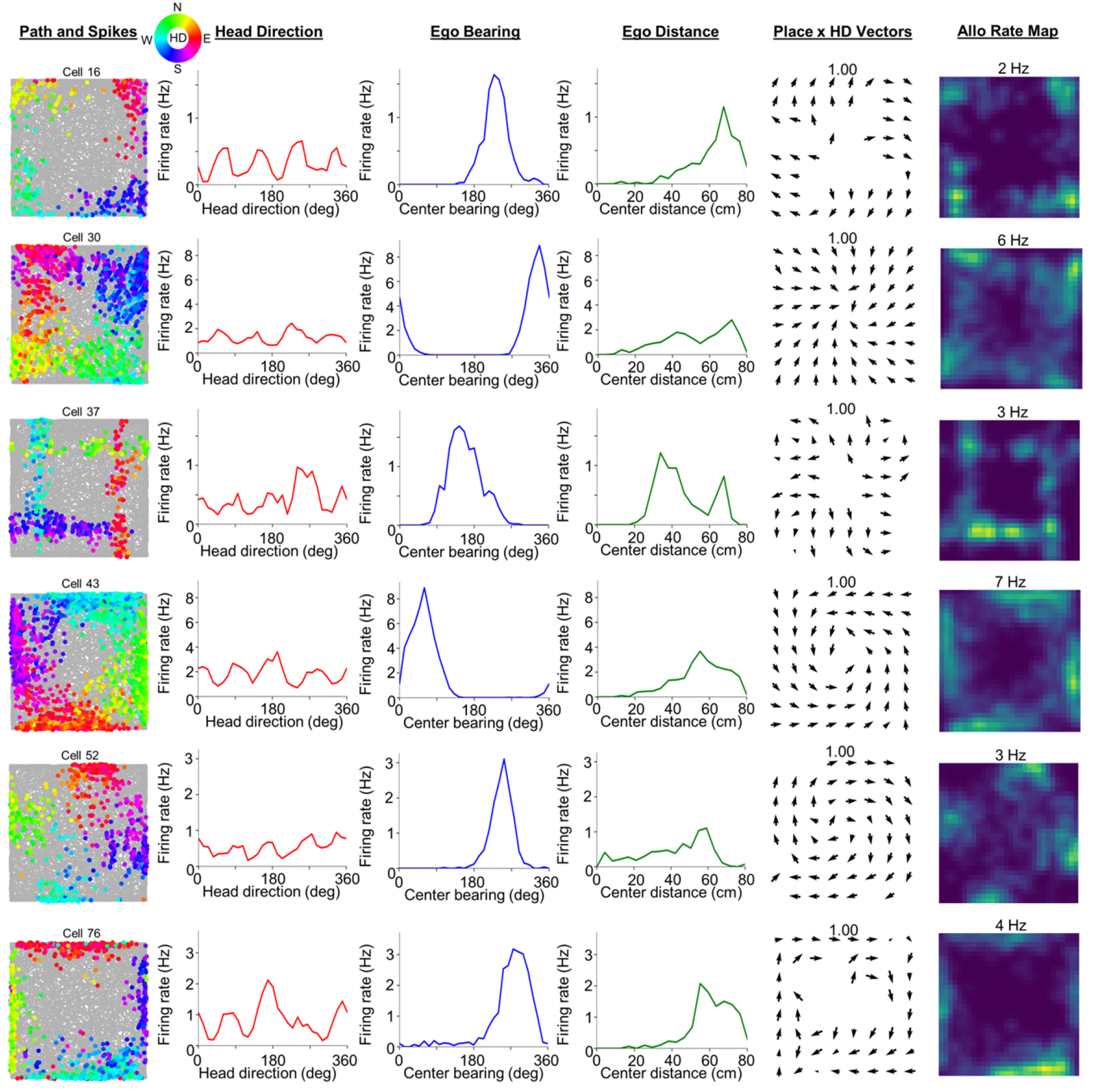
Modeled V1-RSC egocentric bearing cells. Directional spike plots, tuning curves, place-by-HD vector plots, and allocentric firing rate maps for six example modeled V1-RSC cells with significant egocentric bearing tuning. The number above the place-by-HD vector plot indicates the highest MVL, while the number above the allocentric rate map indicates the cell’s peak firing rate.

RSC EB cells have been shown to be strongly modulated by local geometric features (e.g., flat walls and corners), such that their directional tuning exhibits strong four-fold rotational symmetry in a square environment (Alexander et al., 2020; LaChance and Hasselmo, 2024). In contrast, POR EB cells have been shown to lack this strong four-fold symmetry, implying a more global account of environmental geometry that is less impacted by local features (LaChance and Hasselmo, 2024). We assessed four-fold rotational symmetry among the modeled EB cells across three domains: 1) tuning to four distinct HDs spaced 90° apart, assessed using each cell’s HD tuning curve; 2) a distinct firing pattern associated with each encoded HD, assessed by computing cross-correlations between spatial firing rate maps constructed from epochs where the animal was facing distinct HDs; and 3) placement of distinct firing fields at 90° rotational offsets relative to the environment center, assessed using a GLM (see LaChance and Hasselmo, Methods, 2024). Four-fold symmetry scores were computed for each domain based on an autocorrelation analysis (see Methods 2.4.4). The three scores for each cell were then combined to produce an aggregate score, to provide an overall assessment of symmetrical tuning. We found that V1-RSC cells exhibited higher degrees of rotational symmetry across all three domains than SC-POR cells, as shown in Fig. 8A-C, including the aggregate score (HD tuning: Z = 2.52, P = 0.012; HD x location correlations: Z = 2.49, P = 0.013; rotational symmetry analysis: Z = 4.98, P = 6.41e-7; aggregate scores: Z = 4.51, P = 6.59e-6; Fig. 8D). This finding mirrored results from experimental data (HD tuning: Z = 9.52, P = 1.68e-21; HD x location correlations: Z = 7.17, P = 7.64e-13; rotational symmetry analysis: Z = 6.54, P = 6.25e-11; aggregate scores: Z = 9.88, P = 5.00e-23; Fig. 8E-H). However, the SC-POR EB cells tended to display higher degrees of four-fold symmetry than the experimental POR EB cells, with 29% of SC-POR cells displaying an aggregate score *>* 0 compared to 7% of experimental POR cells. Proportions of EB cells with aggregate scores *>* 0 among V1-RSC cells (58%) and experimental RSC cells (53%) were similar, as shown inFig. 8D, H. Overall, however, the trend of RSC EB cells showing stronger four-fold symmetry than POR EB cells was apparent in both the modeled and experimental data.

**Figure 8.**
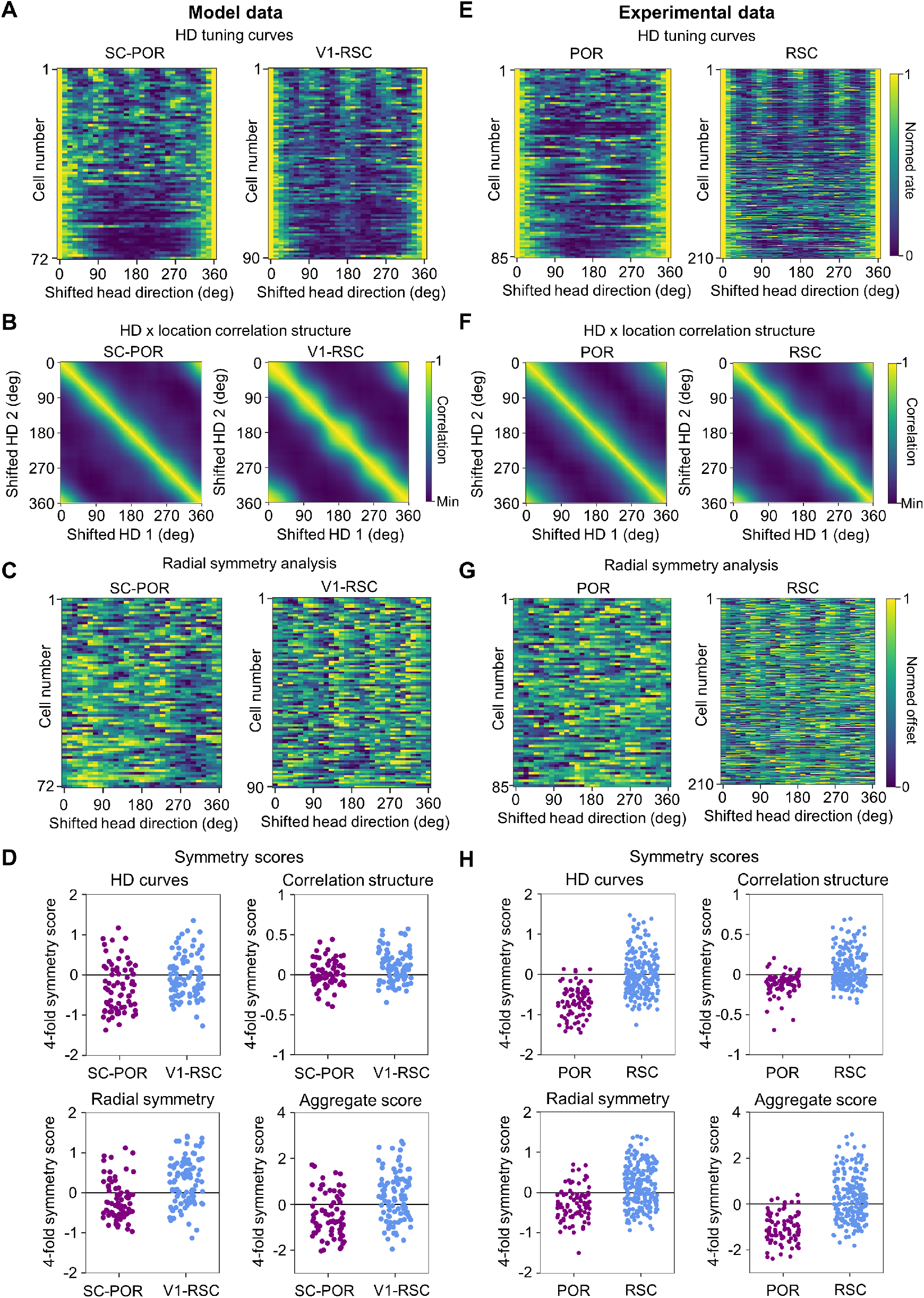
Population coding of environmental symmetry in a square environment. A) Normalized HD tuning curves for all modeled SC-POR and V1-RSC cells with significant EB tuning, shifted for each cell such that the maximum firing rate lies at 0°. Cells are sorted from highest to lowest 4-fold HD symmetry scores. B) Mean HD x location correlation matrix for all SC-POR and V1-RSC EB cells. C) Normalized detrended GLM-derived rotation functions for all SC-POR and V1-RSC EB cells. Cells are sorted from highest to lowest 4-fold radial symmetry scores. D) 4-fold symmetry scores for all modeled EB cells derived from: top left, HD tuning curves; top right, HD x location correlation matrices; bottom left, GLM-derived rotation functions; bottom right, aggregate based on summation of individual symmetry scores. E-H) Same as (A-D) but for experimental neurons recorded from POR and RSC.

In a further test of local vs. global geometric tuning among EB cells, it has been shown that transforming the square environment into an L-shaped environment reveals stronger tuning to local boundary geometry among RSC cells and global boundary geometry among POR cells (LaChance and Hasselmo, 2024). To compare these results to our model cells, we simulated SC-POR and V1-RSC cell firing in a 1.2 × 1.2 m L-shaped environment, shown in Fig. 9A, B, and used a GLM (LaChance and Taube, 2023; LaChance and Hasselmo, 2024) to assess if the cells were more strongly tuned to local or global geometric features (e.g., local boundaries vs. the average location of all boundaries; Fig. 9C). The local and global models were compared using a Globality Index (GI), with higher values indicating a more global geometric signal (see Methods 2.4.3). Comparison between the square and L-shaped environments revealed that SC-POR cells shifted toward a global account of environmental geometry (W = 651, P = 1.99e-4), while V1-RSC cells were better fit by a local account of environmental geometry (W = 1310, P = 3.00e-3; Fig. 9D). This finding mirrored the experimental data, which exhibited a similar distinction between POR and RSC cells in the encoding of local vs. global geometry (POR: W = 289, P = 5.33e-3; RSC: W = 2986, P = 7.84e-13; Fig. 9D).

**Figure 9.**
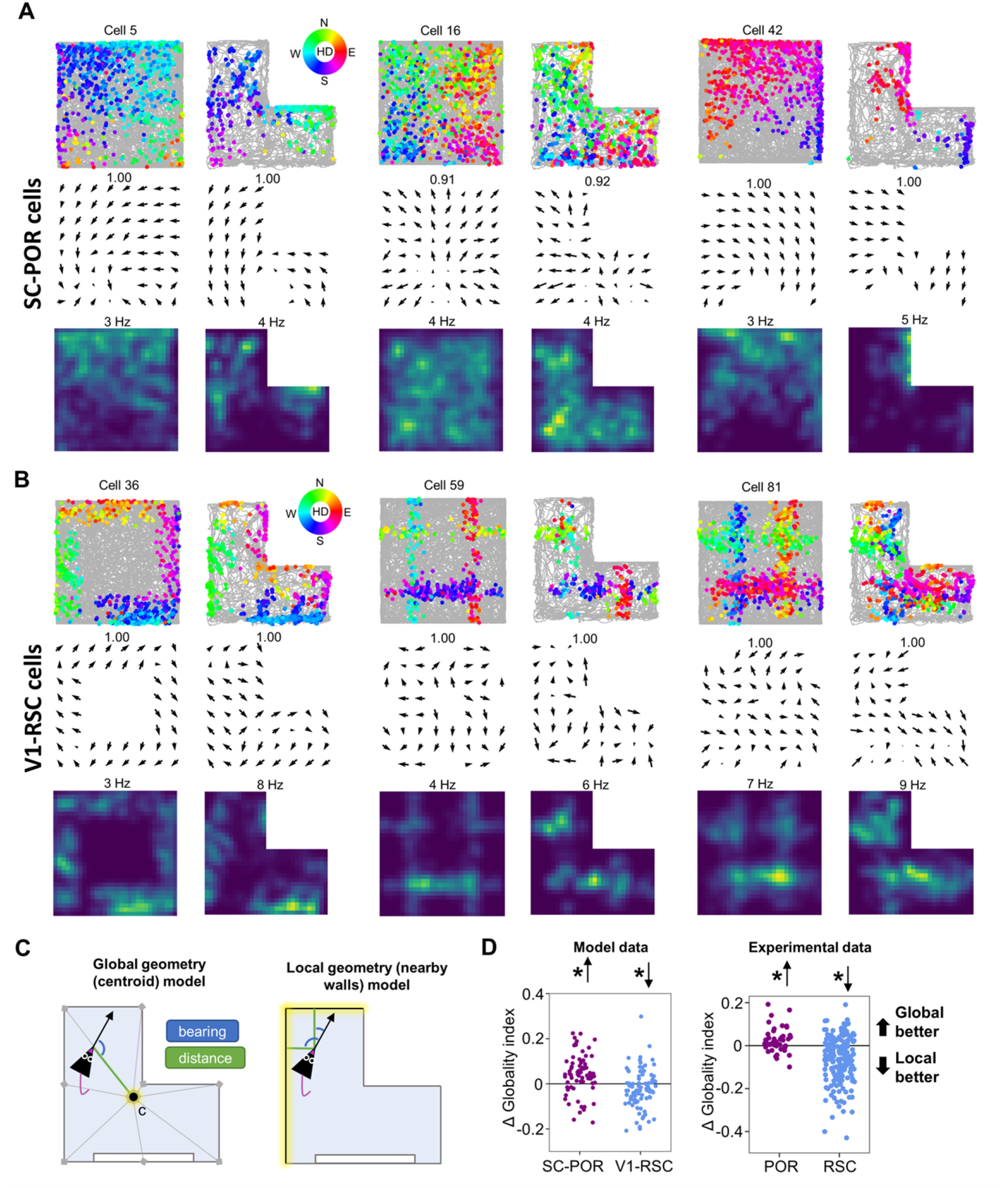
Local vs. global coding of environmental geometry. A) Directional spike plots, place-by-HD vector plots, and allocentric firing rate maps for three example SC-POR cells simulated in both square and L-shaped environments. The number above the place-by-HD vector plot indicates the highest MVL, while the number above the allocentric rate map indicates the cell’s peak firing rate. B) Same as (A) but for three example V1-RSC cells. C) Schematic illustration of the models used to compare local vs. global encoding of environmental geometry. D) Change in globality index between square and L-shaped environments for: left, modeled cells; right, experimental cells. Note that in both modeled and experimental datasets, POR cells tended toward global geometry encoding while RSC cells tended toward local geometry encoding.

### 3.2 Firing properties of model and experimental head direction cells

In addition to egocentric bearing tuning, a large number of SC-POR and V1-RSC cells exhibited apparent tuning to allocentric HD, as shown in Fig. 3A. As the cue card (cue A) along the south wall of the environment provided the only allocentric orienting cue, we hypothesized that this HD tuning was related to visual processing of the cue card. Both POR and RSC HD cells have been shown to be significantly modulated by the presence of similar visual landmarks (Jacob et al., 2017; Zhang et al., 2022; LaChance et al., 2022; LaChance and Hasselmo, 2024). Indeed, the preferred HDs of model HD cells appeared to be biased toward the direction of the cue card in both SC-POR and V1-RSC populations (270°; Fig. 10A). To test whether the apparent HD signal was driven by the presence of the cue card, we simulated SC-POR and V1-RSC cell firing in a square environment with two cue cards placed on opposite walls (cue A and cue B; Fig. 10B). Both SC-POR and V1-RSC HD cells fired in two opposite directions in this condition, assessed using a bidirectionality index (LaChance et al., 2022; SC-POR: W = 0, P = 1.63e-10; V1-RSC: W = 0, P = 7.58e-12; Fig. 10C-E). Further, the two cues were represented with relatively equal firing rates, assessed using a modulation index (LaChance et al., 2022), although SC-POR cells showed slightly higher modulation by cue A than cue B (SC-POR: W = 2638, P = 0.038; V1-RSC: W = 3019, P = 0.30; Fig. 10I). While experimental POR and RSC HD cells also tended to display overall bidirectional firing in this condition (POR: W = 1, P = 2.33e-10; RSC: W = 111, P = 1.41e-4; Fig. 10F-H), firing to the more familiar cue A was much more robust than firing to the more novel cue B (POR: W = 49, P = 3.53e-5; RSC: W = 3, P = 7.28e-11; Fig. 10J), suggesting that other non-visual signals have influence over the POR and RSC cells to cause them to represent cue A more strongly, unlike SC-POR and V1-RSC cells which respond purely to visual properties of the environment.

**Figure 10.**
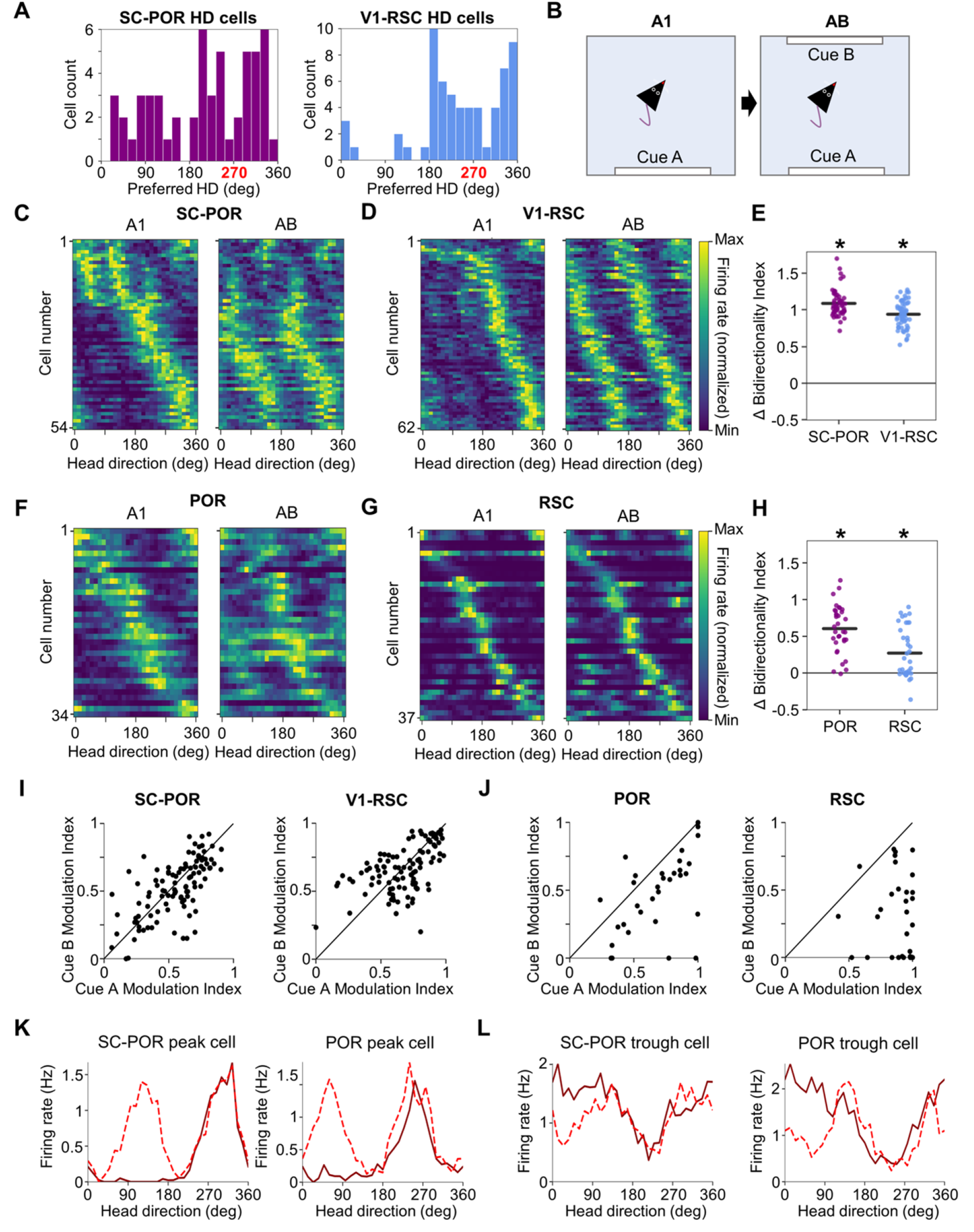
Coding of bidirectional symmetry by HD cells. A) Distribution of preferred HDs for SC-POR and V1-RSC HD cells. B) Schematic showing experimental design for the cue duplication experiment. C) Normalized tuning curves for SC-POR HD cells simulated in both A1 and AB sessions. D) Same as (C) but for V1-RSC HD cells. E) Change in bidirectionality index between the A1 and AB sessions for SC-POR and V1-RSC HD cells. F-H) Same as (C-E) but for experimental neurons recorded from POR and RSC. I) Comparison of the amount of firing rate modulation attributed to cue A vs. cue B in the AB session for SC-POR and V1-RSC HD cells. J) Same as (I) but for experimentally recorded POR and RSC cells. K) Example tuning curves for an SC-POR HD cell and POR HD cell that showed a duplication of their tuning curve peak in the AB session. L) Same as (K) but for cells that duplicated their tuning curve trough in the AB session.

One striking property of the simulated SC-POR HD cells is that they appeared to comprise two separate populations: one that fired most strongly in the general direction of the cue (preferred direction *<* 180°) and contained a sharp peak in its tuning curve; and one that appeared to be inhibited in the direction the cue (preferred direction *>* 180°) and contained a sharp trough in its tuning curve, as shown in Fig. 10C, F. Indeed, in the two cue condition, ‘peak cells’ tended to adopt a second peak 180° opposite the first (Fig. 10K), while ‘trough cells’ adopted a second trough (Fig. 10L). These properties are highly similar to the ‘peak cells’ and ‘trough cells’ found in experimental POR data (LaChance et al., 2022; LaChance and Hasselmo, 2024). As with the experimental RSC data, V1-RSC cells largely lacked trough-related firing (Fig. 10D, G), and in fact almost exclusively exhibited tuning curve peaks in the general direction of the cue card (Fig. 10A, D). While this strong concentration of preferred HDs toward the cue card among V1-RSC cells is unlike experimentally recorded RSC HD cells (Fig. 10G), this property of the model cells may provide insight into how visual signals might interact with HD representations in RSC.

## 4 Discussion

### 4.1 Role of different visual pathways

In this study, we build a learning model based on the visual pathway from SC to POR and demonstrate that diverse spatial properties such as HD tuning and egocentric tuning in POR can be learnt from visual input that processes motion. By comparing our previously designed V1-RSC learning model (Lian et al., 2023) for the area of the RSC, we show that experimentally discovered distinct spatial properties in RSC and POR (LaChance and Hasselmo, 2024) can be largely accounted for by our models, V1-RSC and SC-POR. Note that the V1-RSC model and SC-POR model only differ in their visual inputs; namely that V1 processing represents static feature selectivity similar to simple and complex cells in V1, and SC processing represents visual motion selectivity similar to neurons in SC. Therefore, we conclude that these distinct properties in RSC and POR may originate from the upstream input of these disparate V1 and SC visual pathways. Given that both POR and RSC project to and receive feedback from other areas, including the entorhinal cortex and hippocampus, it is possible that both visual pathways contribute to the brain’s internal map of the external environment. Furthermore, feature processing in the V1 pathway may contribute more to coding of local landmarks, while motion processing in the SC pathway may contribute more to coding of the global environment.

### 4.2 Comparison between model and experimental data

Our model can account for diverse spatial properties in both RSC and POR properties, but there are also discrepancies between model and experimental data in some aspects, such as the percentage of different cell types (Fig. 3), symmetry scores of POR (Fig. 8 DH) and bidirectional symmetry (Fig. 10). One factor might be that the model assumes that the head direction of the virtual rat is aligned with movement direction, whereas there will be some jitter between head and movement directions in experimental studies. More importantly, only visual input is provide to the model. Moreover, to account for distinct spatial properties in RSC and POR, we use two disparate vision inputs. Though model data can capture diverse spatial properties in POR or RSC and the major difference between them, upstream input in the model is much less complicated compared with the input neurons in POR and RSC receive in real neural circuits. As more experimental studies reveal the upstream input these cortices receive, the model could be improved to incorporate these inputs and, possibly, more closely match the experimental data.

### 4.3 Underlying learning principles of these cortical spatial representations

Both SC-POR and V1-RSC learning models are based on the principle of sparse coding (Olshausen and Field, 1996, 1997) that has been demonstrated to account for the emergence of other spatial cells in the brain’s navigational system (Lian and Burkitt, 2021, 2022; Lian et al., 2023). However, this does not suggest that sparse coding is the only principle that can contribute to learning spatial cells from visual input, especially given its relationship with other neural organizing principles such as predictive coding and divisive normalization (Lian and Burkitt, 2024), and potential other principles.

### 4.4 Implications of the study for research on spatial neurons

The results of this study have significant implications for our understanding of various spatial cell types found throughout the brain. Neurons that respond to environmental geometry in an egocentric reference frame have been reported in a variety of regions including POR (Gofman et al., 2019; LaChance et al., 2019), RSC (Alexander et al., 2020; van Wijngaarden et al., 2020), lateral entorhinal cortex (Wang et al., 2018), dorsal presubiculum (Peyrache et al., 2017), and dorsal striatum (Hinman et al., 2019), though it remains unclear how the egocentric response properties in these brain regions may differ from each other. Consideration of the specific visual inputs to these regions may provide insight into the mechanisms behind their egocentric firing (e.g., optic flow vs visual feature processing), as well as applying the rotational symmetry and local vs. global analyses outlined here and in a previous study (LaChance and Hasselmo, 2024). It is worth considering that POR and RSC are also reciprocally connected (Burwell and Amaral, 1998a; Agster and Burwell, 2009) and share connections with all of the regions listed above (Sugar et al., 2011; Monko and Heilbronner, 2021; Estela-Pro and Burwell, 2022), so the EB cells in each brain area may show heterogeneity or mixed response properties given their varied inputs. It has been shown previously that even POR EB cells can show heterogeneity in their responses to local vs. global aspects of environmental geometry (LaChance and Taube, 2023) despite the overall population being significantly global-shifted when compared to the more local-shifted RSC EB cells (LaChance and Hasselmo, 2024), so individual cells in each region likely fall along a continuum between visual motion and visual feature processing.

One particularly notable aspect of the simulated dataset is its generation of HD cells that overall match the distinct firing properties of empirically recorded HD cells in POR and RSC (LaChance et al., 2022). Many HD cells in both POR (LaChance et al., 2022; LaChance and Hasselmo, 2024) and RSC (Jacob et al., 2017; Zhang et al., 2022; Sit and Goard, 2023; LaChance and Hasselmo, 2024) (but see Lozano et al. (2017)) have been shown to be capable of firing along two opposite preferred directions when a visual landmark is duplicated along two opposite walls of an environment, an effect captured by both SC-POR and V1-RSC models presented here. However, three important properties of the simulated cells may provide special insight into the integration of visual landmarks into the HD system. First, much like the empirical POR data, only the SC-POR model produced both peak and trough cells (i.e., HD and anti-HD cells), whereas the V1-RSC model produced only peak cells, suggesting that optic flow processing may be especially suited for producing the kind of dichotomous (toward landmark vs. away from landmark) firing preferences observed among HD cells in POR. Second, the secondary peak or trough adopted by the SC-POR and V1-RSC cells was generally the same size as the original peak or trough, unlike the empirical data where the original peak or trough was almost always larger. This effect in the empirical data may be due to input from vestibular-based ‘classic’ HD cells (Taube et al., 1990a; Yoder and Taube, 2014) which continue to fire in a single direction (the ‘true’ allocentric direction) despite bidirectional symmetry of visual landmarks (LaChance et al., 2022) and which were not simulated in the current study. Third, the V1-RSC model almost entirely produced cells with preferred directions oriented toward the cue card, whereas empirical RSC HD cells have uniformly distributed preferred directions (Jacob et al., 2017; Zhang et al., 2022; Sit and Goard, 2023; LaChance and Hasselmo, 2024). As with the previous point, this discrepancy could likely be corrected by incorporating inputs from ‘classic’ HD cells with a uniform distribution of preferred directions, which can be bound to the external world by the visually-based ‘HD’ cells produced by the V1-RSC model. Thus, the exclusive presence of landmark-directed cells in the V1-RSC model hints at how visual landmark processing in RSC may differ from and integrate with the ‘classic’ HD signal to bind it to specific environmental features (also see Bicanski and Burgess (2016); Page and Jeffery (2018); Yan et al. (2021)).

These results also suggest ways in which POR and RSC neurons may differentially impact allocentric spatial cell firing in downstream regions. Notably, both POR and RSC provide strong inputs to the hippocampal formation, including direct projections to the entorhinal cortex (Wyass and Van Groen, 1992; Burwell and Amaral, 1998b; Koganezawa et al., 2015; Doan et al., 2019) and subiculum (Wyass and Van Groen, 1992; Naber et al., 2001). The medial subdivision of the entorhinal cortex (MEC) in particular contains allocentric grid cells (Hafting et al., 2005), border cells (Solstad et al., 2008), and object vector cells (Høydal et al., 2019), whereas the subiculum contains allocentric boundary vector cells (Lever et al., 2009), corner cells (Sun et al., 2024), and geometry-agnostic place cells (Sharp, 2006), all of which are likely to be informed by the egocentric visual representations in upstream POR and RSC. Future physiology studies could investigate how each of these allocentric spatial cell types may be differentially impacted by visual motion processing in POR or visual feature processing in RSC. For example, inactivating POR may disrupt the MEC grid cell global firing pattern but not affect the stability of firing fields relative to local boundary features, whereas inactivating RSC may cause unstable firing near boundary features despite maintenance of the overall structure of the grid pattern. Both optic flow (Raudies et al., 2012; Raudies and Hasselmo, 2012) and visual features (Alexander et al., 2023) have been proposed to shape both grid cell and boundary vector cell firing. Other spatial cell types should be considered in terms of these visual information streams and their relative contributions to cell firing, as well as neurally plausible transformations that must take place to integrate the specific egocentric representations in POR and RSC into an allocentric reference frame downstream in the hippocampal formation.

## Acknowledgements

This work received funding from the Australian Government, via grant AUS-MURIB000001 associated with ONR MURI grant N00014-19-1-2571. This research was also supported by NIH NINDS K99 NS119665, NIMH R01 MH120073; Office of Naval Research MURI grant N00014-16-1-2832; Office of Naval Research MURI N00014-19-1-2571; and Office of Naval Research DURIP N00014-17-1-2304.

## Notes

### Competing Interest Statement

The authors have declared no competing interest.

